# Rapid, Growth Factor-Reduced Differentiation of Functional Neurons from hiPSCs

**DOI:** 10.64898/2025.12.03.692194

**Authors:** Natalie A Parker, Oluwadamilola E Kolawole, Zhixin Liao, Farsin S Syed, Kara E Moquin, Nisha R Iyer

## Abstract

**Introduction:** Human induced pluripotent stem cells (hiPSCs) can be rapidly converted into neurons via NGN2 overexpression, but many protocols require costly reagents during the initial induction phase that may limit adoption by labs without routine neuronal differenitation experience. We developed a simplified, low-cost protocol using a tetracyline-inducible (TET-on) NGN2 system in minimal media to generate cortical neurons in as little as 6 days.

**Methods:** KOLF2.1J hiPSCs were stably transfected with a TET-on NGN2 cassette using the nonviral PiggyBac system and induced with doxycycline in Essential 6 media. The impact of adding the Notch inhibitor, DAPT, during doxycycline induction to enhance neurogenesis to was evaluated with immunocytochemistry (ICC) and RT-PCR. Following induction neurons were matured and characterized with ICC for mature neuronal markers and by multielectrode array recordings for functional network activity.

**Results:** DAPT markedly improved conversion efficiency, reducing non-neuronal cells and increasing pan-neuronal TUJ1 expression. Resulting neurons expressed cortical markers and matured into functional glutamatergic neurons. MEA recordings showed spontaneous activity by day 14 and synchronous network firing by day 35. Secondary PB transfection enabled Td-Tomato labelling of KOLF2.1J:pB-TO-NGN2 hiPSCs, allowing 24-hour live imaging of neurite outgrowth.

**Conclusion:** This streamlined, growth-factor-free workflow provides an accessible route for generating functional neurons from patient-derived hiPSCs, including in labs with limited hiPSC or neuronal culture experience.

## Introduction

Human pluripotent stem cells (hPSCs) are a critical tool for modelling human neurodevelopment, studying neurological diseases and therapeutic discovery (1,2), and the growing use of patient-derived induced pluripotent stem cells (hiPSCs) holds promise for personalized medicine (3). Yet despite the availability of hiPSC-derived neurons, many labs still rely on primary rodent neurons or immortalized cell lines, largely because differentiation methods remain technically demanding and expensive to adopt. Classic directed differentiation approaches apply small molecules and growth factors to guide neuronal fate by mimicking development, but non-experts may be hampered by long differentiation timescales (weeks to months), costly growth factors, poor reproducibility between hiPSC lines (4–6). Direct reprogramming by tetracycline inducible (TET-on) overexpression of master regulator NGN2 has emerged as a faster (days) and more reproducible approach to generate highly enriched hPSC-derived neuron cultures (7–13). (7,11,13Due to its scalability and culture consistency, TET-on NGN2 reprogramming is being applied to large hiPSC collections, or “villages,” to model donor-specific genetic variability in disease, thus advancing personalized patient models for mechanistic studies and drug screening (14,15).

Despite these comparative advantages, many published NGN2-based methods still apply growth factors and media supplements during the DOX induction phase, introducing similar cost and reproducibility concerns as for directed differentiation (7–13). Dual-SMAD inhibition is commonly applied to promote neuronal lineage commitment, while additional signaling factors, such as Wnt agonists, retinoic acid, or sonic hedgehog, are used to guide NGN2 reprogramming toward specific neuronal subtypes (10). For example, these patterning cues have been employed to generate excitatory glutamatergic (8,13), dopaminergic (16), or spinal motor neurons (14) or are provided through commercially available differentiation kits. NGN2 induction efficiency and scalability are also limited by the transgene delivery method. Lentiviral transduction (9–11,17) or CRISPR knock-in (7) are common, but can be time-consuming, expensive, and carry non-trivial insertional mutagenesis risk (18). In addition, the need for BSL-2+ facilities and specialized gene editing or viral expertise may limit adoption by under resourced labs or emerging scientists.

Our goal was to develop a simple, growth factor-reduced NGN2 differentiation method using piggyBac (PB) transposon tools recently developed by the iPSC Neurodegenerative Disease Initiative (iNDI) (19,20). The PB system is a high-efficiency “cut-and-paste” gene delivery method whereby the PB transposase integrates the desired gene cassette at TTAA sites (21,22). Unlike Sleeping Beauty or Tol2, PB exhibits high transposition activity across species and supports precise, scar free excision (21,23,22). Its capacity for large payloads and repeated transfections makes it well suited for engineering hPSC lines with inducible transgenes (16,17), including TET-on NGN2 for neurons (19). We rationalized that we could simplify differentiation of PB-edited polyclonal hiPSCs because media additives are often inherited from direct differentiation protocols to support neuroepithelial specification. However, neuroepithelial induction can be achieved with Essential 6 (E6) media alone (24) and its simple six-component composition enables in-house media production.

Here we combine simple E6-based neural induction with PB–TO-NGN2, and show that with modest optimization this yields high-efficiency cortical-like neurons. Upon exposure to neurotrophins, these mature rapidly into functional glutamatergic neurons with synapses and networked electrophysiology. Finally, we show that secondary PB transfection with fluorescent reporters enables live imaging, providing a practical on-ramp for labs to transition from rodent models to human cells for *in vitro* applications.

## Methods

### Stem Cell Maintenance

Experiments were performed using KOLF2.1J hiPSCs (JAX; passage range 10-50), WA09 (H9) hESCs (WiCell; passage range 31-37), or WA01 (H1) hESCs (WiCell; passage range 40-45) cultured at 37°C with 5% CO_2_. hPSCs were maintained in Essential 8 (E8) medium (ThermoFisher Scientific) on tissue culture–treated plates coated with growth factor-reduced Matrigel (Corning). Cells were passaged when they reached 70–90% confluency using Versene (24,25) (ThermoFisher Scientific) into E8 media with Chroman 1 (50 nM), Emricasan (5 µM), Polyamine (1X), and Trans-ISRIB (700 nM) (CEPT) (26); E8 media was changed daily. Cell lines were tested monthly for mycoplasma with MycoAlert (Lonza).

### Generation of Stable PiggyBac Insertion Cell Lines

To generate polyclonal cell lines with inducible NGN2 expression, hPSCs were transfected with the PB-TO-hNGN2 vector (gift from the iPSC Neurodegenerative Disease Initiative (iNDI) & Michael Ward; Addgene plasmid #172115) when ∼50–70% confluent as previously described (27). Briefly, media was first replaced with E8 +CEPT. A DNA mixture containing 0.75 µg PiggyBac transposase and 2.25 µg transposon plasmid was diluted in Opti-MEM to 100 µL and combined with 5 µL Lipofectamine Stem in 95 µL Opti-MEM. After a 20-minute incubation, the 200 µL mix was added dropwise to each well and cells were incubated overnight. Media was changed daily with E8+CEPT until cultures reached 70–90% confluency. Cells were then passaged 1:6 with Versene and replated on Matrigel in E8+CEPT. 24 hours post-passage, transfection efficiency was determined by manually counting the number of BFP^+^ cells out of total cells from 3 fields of view in a transfected well. Media was then replaced with selection medium (E8+CEPT + 8 µg/mL puromycin) and refreshed daily. Cultures were monitored for BFP until ∼100% positivity (typically days 8–10), after which CEPT and puromycin were withdrawn, and lines were maintained under standard hPSC conditions.

For secondary transfection of KOLF2.1J:pB-TO-NGN2 hiPSCs, cells were transfected with PB-CAG-tdTomato (Addgene #1331569) and PBCAG-eGFP (Addgene #40973) using the same procedure, passaged into E8+CEPT and subsequently maintained under standard conditions.

### NGN2 Induction and Maturation

To initiate differentiation, KOLF2.1J:pB-TO-NGN2 hiPSCs were singularized with Accutase (StemCell Technologies) and seeded at 50,000 cells/cm^2^ in E8 +CEPT on Matrigel-coated plates. After 24 hours, cells were washed with 1× PBS and switch to E6 media (24) (ThermoFisher Scientific) containing 2 µg/mL doxycycline (28) (Millipore Sigma) with or without 10 µM DAPT (Tocris Bioscience). 80% media volume was replenished daily for six days. Beginning on Day 3, Laminin (1 ng/mL; ThermoFisher Scientific) was added to support neuronal adhesion.

For maturation, neurons were singularized and replated on Matrigel-coated plates at 10,000 cells/cm^2^ and cultured on maturation media Neurobasal Plus (ThermoFisher Scientific) with B27 Plus (1X; ThermoFisher Scientific), N2 (1X; ThermoFisher Scientific), GDNF (10 ng/mL; PeproTech), BDNF (10 ng/mL; PeproTech), Ascorbic Acid (200 μM; Sigma-Aldrich), cAMP (1 μM; Millipore Sigma), and Laminin (1 ng/mL) (7, 28). Maturation media was replenished with 50% changes every 3-4 days.

### Gene Expression Analysis

Cells were lysed directly in TRIzol reagent (Invitrogen) and RNA was extracted using Phasemaker tubes with chloroform phase separation, isopropanol precipitation, and ethanol washing following manufacturer protocols (ThermoFisher Scientific). Pelleted RNA was resuspended in RNase/DNase-free water, and concentration/purity were assessed using a DeNovix spectrophotometer. cDNA was generated from purified RNA using the SuperScript IV First-Strand Synthesis System (ThermoFisher Scientific) according to the manufacturer’s instructions. For each 20 µL reaction, 1,000 ng of cDNA was diluted to 9 µL in RNase/DNase-free water and combined with 10 µL TaqMan Gene Expression Master Mix and 1 µL TaqMan Gene Expression Assay (ThermoFisher Scientific; Table S1). Reactions were run on a QuantStudio 3 Real-Time PCR System (Applied Biosystems) with the protocol: 50°C for 2 min, 95°C for 2 min, and 40 cycles of 95°C for 15 sec followed by 60°C for 1 min. This assay included three technical replicates per sample and a no-cDNA negative control for each gene to confirm the absence of non-specific amplification. Relative gene expression differences were calculated with the △△C_t_ method, normalized to RPS18 expression.

### Electrophysiology

Electrophysiological activity was recorded on the Maestro Edge MEA system (Axion Biosystems). Differentiated neurons were seeded onto 24-well CytoView MEA plates at 100,000 cells per well. To localize cells over the electrodes (**Fig. 3a**), 400 µL of cell-containing media was pipetted onto the center of each well and allowed to absorb for 30 minutes at 37°C before adding the remaining media. 50% media changes occurred at least 24 hours before recording to minimize artifacts. Spontaneous activity was recorded for 5 minutes at each timepoint. After spontaneous recordings, neurons were recorded while electrically stimulated 5 times for 500 µs at 600mV and 500µA. Raster plots were generated using AxIS Neural Metric Software (Axion Biosystems) and adjusted firing rate and network synchrony were quantified using RStudio.

On day 34 of maturation, neurons were treated with 20 µM CNQX to inhibit AMPA receptors (30,31),with spontaneous activity measured before and during treatment, after which CNQX was removed by one full media change followed by three consecutive 50% media changes.

Spontaneous activity was recorded again 24 hours after the wash-out. Adjusted mean firing rate and synchrony index were quantified over the following intervals: the 5 minutes immediately before treatment, the 15-minute treatment period divided into 5-minute bins, and a 5-minute window collected 24 hours after washout (**Fig. 3e**).

### Immunocytochemistry (ICC) and Quantification

Prior to immunocytochemistry (ICC) and quantification, cultures were singularized with Accutase and replated at 1,000 cells/cm^2^ on glass-bottom wells for 12 hours. Cells were fixed for 20 minutes in 4% paraformaldehyde mixed 1:1 with culture media, then washed three times with TBS. Blocking was performed for 1 h at room temperature in TBS with 0.3% Triton X-100 and 5% Normal Donkey Serum (TBS-DT). Primary antibodies (Table S2) diluted in TBS-DT were applied overnight at 4 °C. After three washes in TBS with 0.3% Triton X-100, AlexaFluor secondary antibodies (1:500 in TBS-DT) were added for 1 h at room temperature. Samples were washed twice in TBS, stained with DAPI (1:2000, 10 minutes), washed again, and mounted in Gold Antifade. Imaging was performed on an ImageXpress Micro Confocal (Molecular Devices) and processed in ImageJ. TUJ1+ cells were quantified as a fraction of DAPI+ nuclei from five randomly selected fields per replicate; fields without cells were replaced.

### Live Imaging

For live imaging, tdTomato-expressing cells were replated at 10,000 cells/cm^2^ on glass bottom 96-well plates and maintained in Neurobasal Plus supplemented with B27 Plus and N2). Plates were sealed on the ImageXpress Micro Confocal stage and maintained at 37 °C and 5% CO_2_. Images were collected every 5 minutes for 24 hours, and images were processed in ImageJ.

### Statistical Analysis

All statistical analyses were performed in R Studio with at least three biological replicates per experiment unless noted. Student’s t-test was used for paired comparisons, one-way ANOVA for gene expression, and repeated measures ANOVA for CNQX treatment time courses, with Tukey’s HSD for post hoc analysis. Results are reported as mean ± standard deviation, with significance defined as *p<0.05, **p<0.01, ***p<0.001.

## Results

### Generation and Induction of NGN2 Neurons in Minimal Media Conditions

To achieve stable, scalable transgene delivery across diverse hPSC lines, we used PiggyBac-mediated insertion of a TET-ON NGN2 cassette containing puromycin resistance gene and a BFP reporter (19,21,22). This approach achieved 39.1% transfection efficiency in KOLF2.1J cells, and puromycin selection yielded a homogeneous BFP+ population within 8 days (**Fig. 1a**). To demonstrate utility across multiple hPSC lines, we also transfected H1 and H9 hESCs; while efficiencies were lower compared to KOLF2.1J hiPSCs (15.2% and 15.3%), all cultures were 100% BFP+ after puromycin selection (**Fig. 1a**). While qRT-PCR showed KOLF2.1J:pB-TO-NGN2 hiPSCs upregulated *NGN2* compared to parental controls (**Supplementary Fig. 1a**), indicative of leakiness of the transgene, stem cell cultures uniformly expressed OCT3/4 for at least 40 passages (**Fig. 1b**), demonstrating pluripotency.

**Figure 1.**
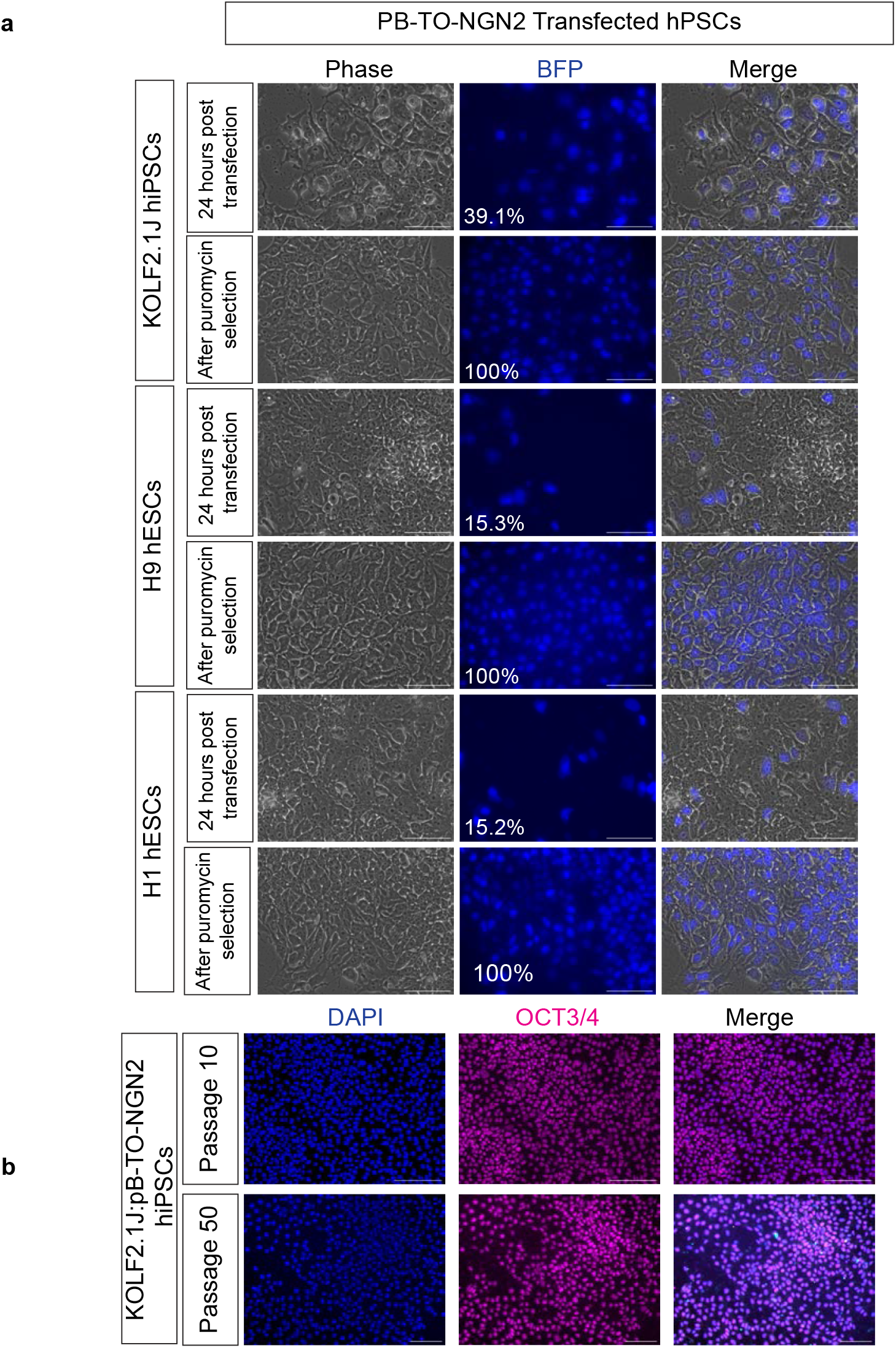
Transfection of PB-TO-NGN2 into multiple hPSC lines. **(a)** BFP+ reporter expression in in KOLF2.1J, H9, and H1 hPSCs 24 hours after transfection of PB-TO-hNGN2 and following puromycin selection. Scale=75 μm **(b)** Immunostaining of Passage 10 and Passage 50 KOLF2.1J::pB-TO-NGN2 iPSCs to confirm pluripotency with DAPI (nuclei; blue) and OCT3/4 (pluripotency; magenta). Scale=150 μm

We next tested whether minimal-media DOX induction could drive robust NGN2-mediated neuronal conversion comparable to other NGN2 methods (7,9). Since Notch inhibition enhances neurogenesis (32), we also tested whether adding γ-secretase inhibitor DAPT could accelerate or improved differentiation (**Fig. 2a**). Clear neurites emerged under DOX conditions by day 3 (**Supplementary Fig. 2a**), and after six days, ICC showed a greater proportion of TUJ1^+^ neurons and reduction of PAX6^+^ progenitors relative to E6 controls (**Fig. 2b**), demonstrating doxycycline-dependent neuronal commitment. qRT-PCR confirmed upregulation of *NGN2* in DOX conditions compared to KOLF2.1J:pB-TO-NGN2 hiPSC controls, though *PAX6* expression remained high in all conditions (**Fig. 2c**). However, neuronal purity in DOX-only conditions was only 65.1% (±5.0), with PAX6^+^ progenitor clusters persisting between TUJ1^+^ regions (**Fig. 2b, d**). Adding the small molecule DAPT significantly improved purity, increasing TUJ1^+^ neurons to 77.6% (±3.6) and reducing the appearance of PAX6^+^ clusters (**Fig. 2b, d**) Throughout the 6 day induction, cultures with DAPT also appeared less dense, consistent with fewer proliferating cells and enhanced differentiation (**Supplementary Figure 2a**). Similar trends were observed in H1 and H9 hESCs, where adding DAPT consistently improved neuronal differentiation compared to DOX alone (**Supplement Figure 2b**). These results demonstrate that the minimal-media NGN2 induction protocol robustly produces postmitotic neurons across multiple hPSC lines.

**Figure 2.**
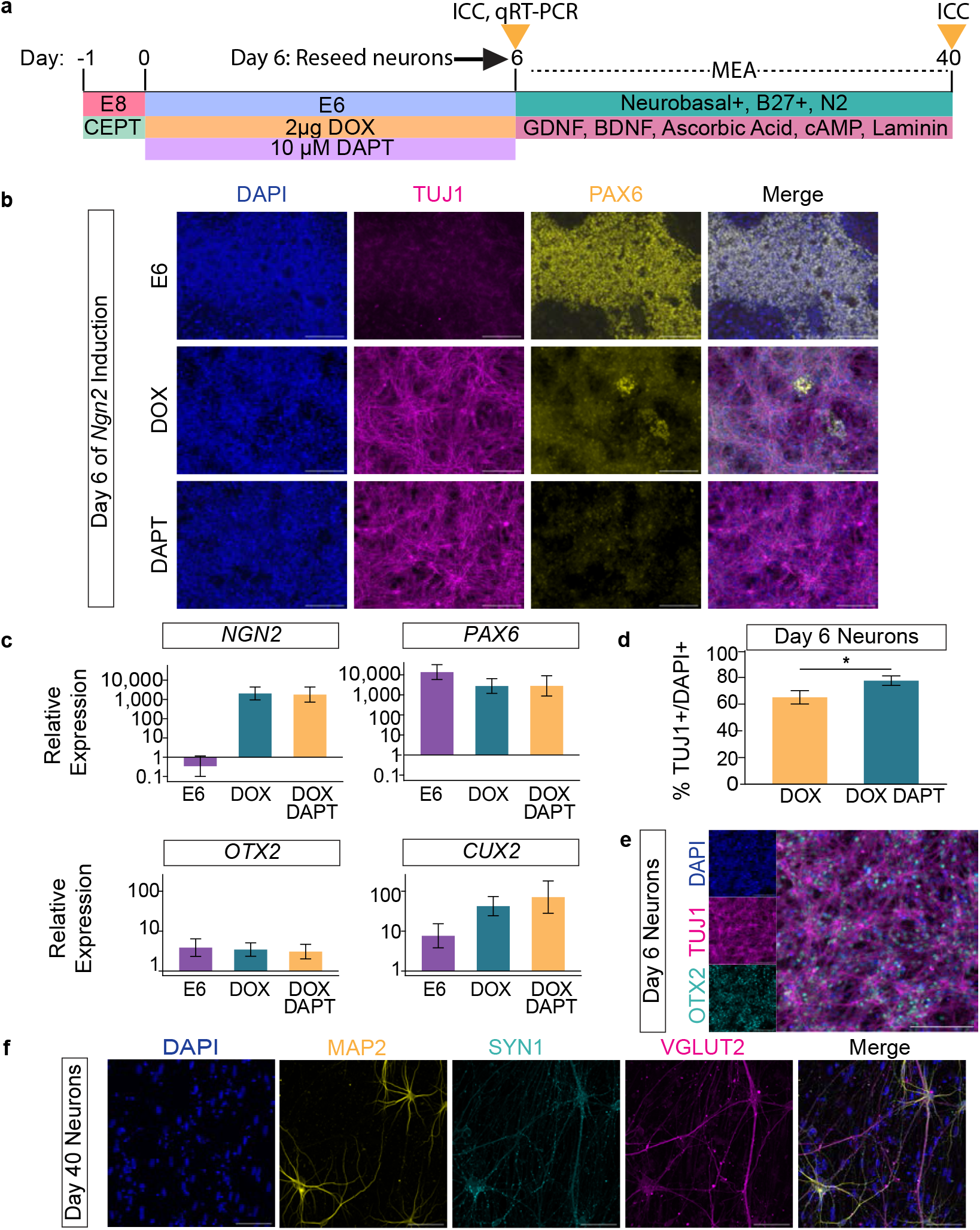
Induction of PB-TO-NGN2 to cortical neurons in minimal media conditions. **(a)** Schematic of induction of neurons in E6, E6+DOX induction, or E6+DOX+DAPT minimal media conditions and subsequent maturation. **(b)** Immunostaining of DAPI (nuclei; blue), PAX6 (neural progenitors; yellow) and TUJ1 (pan-neuronal; magenta) after 6 days of NGN2 induction. Scale=150 μm. **(c)** qRT-PCR of *NGN2* (neuronal), *PAX6* (neuroepithelial), *OTX2* (cortical forebrain), and *CUX2* (cortical) expression at Day 6 of induction. Data shown as relative fold change compared to KOLF2.1J:pB-TO-NGN2 hiPSCs. Error bars shown as standard deviation (n=3) **(d)** Quantification of Tuj1^+^ neurons out of total DAPI^+^ cells in DOX and DOX+DAPT conditions at Day 6 of induction. **(e)** Immunostaining of DAPI (blue), OTX2 (cyan) and TUJ1 (magenta) at Day 6 of induction. Scale=150 μm. **(f)** Immunostaining of MAP2 (mature neurons; yellow), VGLUT2 (glutamatergic neurons; magenta), and SYN1 (pre-synaptic marker, cyan) at Day 40 after maturation. Scale=75 μm.

Because NGN2-induced neurons typically adopt a cortical fate (7,9), we examined expression of cortical markers OTX2, which is present in both cortical progenitors and neurons (34), and CUX2, a transcription factor expressed in post-mitotic forebrain neurons (35). *OTX2* and *CUX2* were expressed in all conditions, but *CUX2* was 10-fold higher in both DOX and DOX/DAPT conditions relative to E6 controls, indicating specification toward post-mitotic forebrain neurons (**Fig. 2c**). When matured in neurotrophic factors, TUJ1^+^/OTX2^+^ cultures (**Fig. 2e)** co-express MAP2, VGLUT2, and SYN1 at Day 40, demonstrating mature glutamatergic fate and formation of synaptic networks (**Fig. 2g, Supplementary Fig. 3**). This is consistent with other NGN2 induction approaches that yield glutamatergic neuronal identities and exhibit a comparable maturation timeline (7,9).

### Functional Maturation of Induced Neurons

We next assessed whether rapidly induced neurons were electrophysiologically active by culturing them on MEAs in maturation media supplemented with neurotrophins (**Fig. 2a, 3a**). By Day 14, neurons exhibited spontaneous firing (**Fig. 3b**). Mean firing rate and network synchrony increased from Day 14 to Day 35, although differences were not statistically significant, likely due to variability across MEA wells (**Fig. 3b, c**). Neurons also responded to electrical stimulation beginning on Day 14 and continuing throughout maturation (**Supplementary Fig. 4**). To confirm excitatory neuron identity (7,9,10), we selectively inhibited AMPA receptors with CNQX (30,31), which reversibly silenced activity, consistent with glutamatergic phenotype (**Fig. 3e-h**). These results show that NGN2-induced neurons rapidly develop functional glutamatergic networks, validating their suitability for downstream applications.

**Figure 3.**
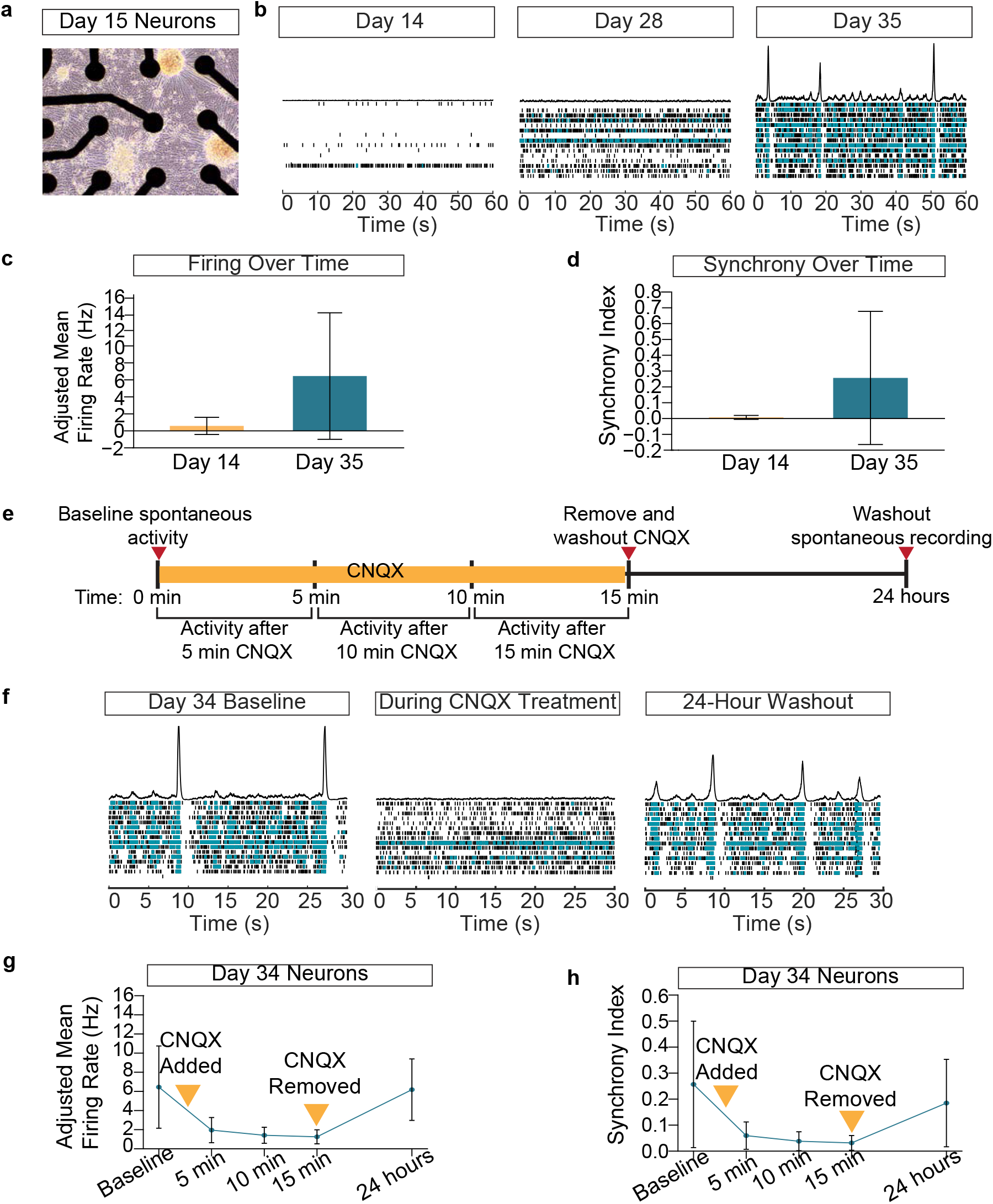
MEA recordings demonstrate NGN2-induced glutamatergic neurons develop spontaneous electrophysiological activity. **(a)** Representative phase contrast image on Day 15 of neuron electrode coverage on an MEA well. **(d)** Representative raster plots from MEA recordings on Days 14, 28, and 35 of culture demonstrating maturation of electrophysiological network function. **(c)** Comparison of adjusted mean firing rate and **(d)** synchrony index between Day 14 and Day 35 (n=3). **(e)** Schematic of MEA recording before, during, and after CNQX treatment. **(f)** Representative raster plots, **(g)** adjusted mean firing rate, and **(h)** synchrony index of spontaneous activity before, during, and after a 24 wash-out of CNQX treatment to inhibit AMPA receptor activity. All error bars represent standard deviation.

### Live Imaging by Secondary PB Transfection

A key application of these neurons is live imaging for high-throughput screening, so we performed a secondary PB transfection using a constitutively active fluorescent reporter, PB-CAG-TdTomato. Notably, secondary transfection into KOLF2.1J:PB-TO-NGN2 hiPSCs produced markedly higher transfection efficiency than transfection into parental KOLF2.1J cells (**Fig. 4a**). We observed the same trend using PB-CAG-eGFP (**Supplementary Fig. 5**). Neither fluorescent PB construct contained a selection cassette, which was acceptable for our purposes because we aimed to sparsely label cells to visualize individual neuronal processes in dense cultures. Timelapse enabled clear visualization of neurite outgrowth over the 24-hour imaging period (**Fig. 4b**), demonstrating that secondary PB transfection supports robust, non-disruptive live tracking of neuronal morphology.

**Figure 4.**
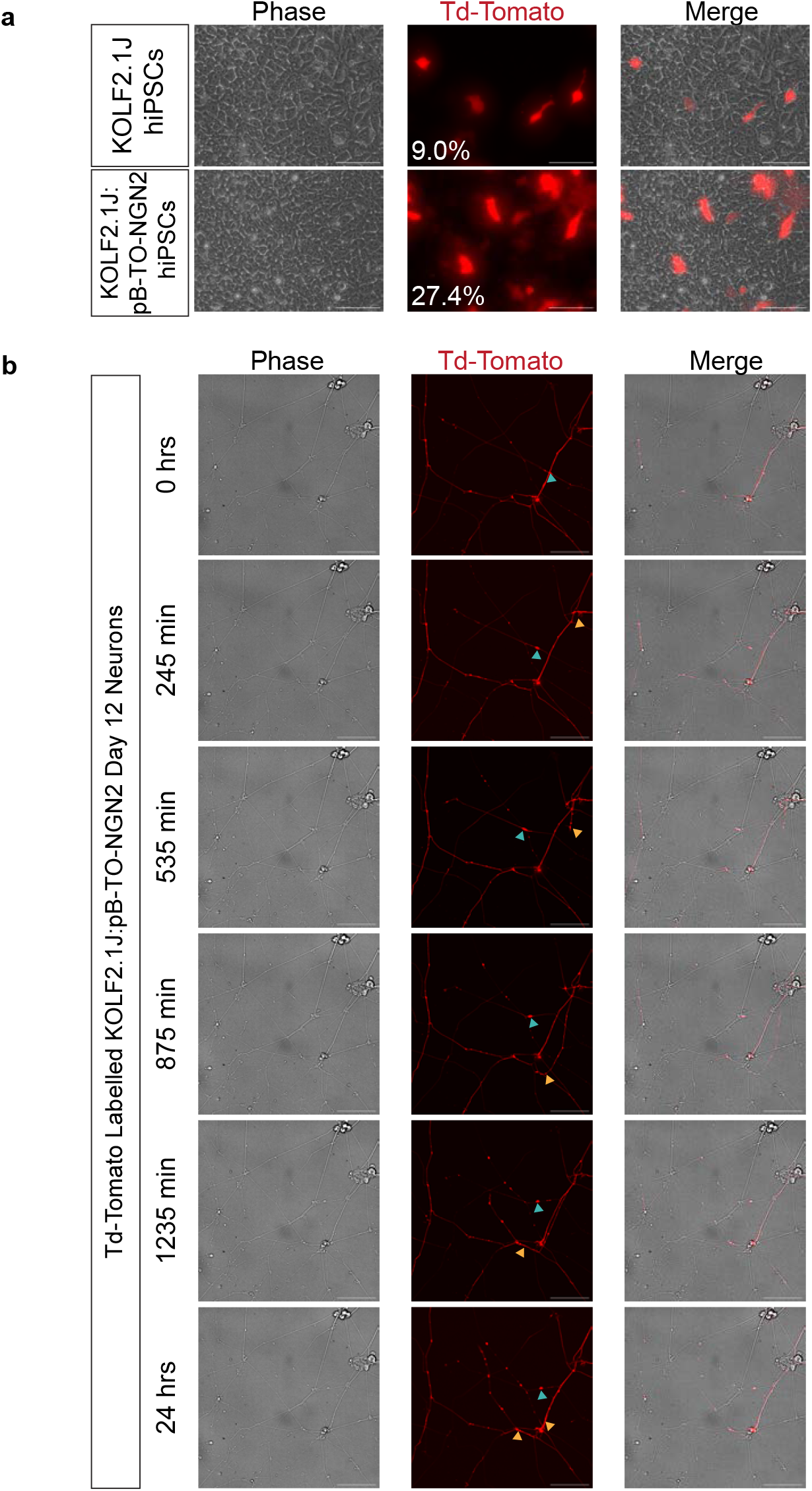
Secondary PiggyBac transfection enables sparse fluorescent labelling for live imaging. **(a)** Transfection efficiency of PB-CAG-Td-Tomato into KOLF2.1J and KOLF2.1J::pB-TO-NGN2 hiPSC lines. **(b)** Representative images over 24 hours of NGN2 induced neurons labelled with Td-Tomato. Arrowheads indicate regions cell body movement or neurite extension. Scale bars=75 μm.

## Discussion

Using a PiggyBac TET-On system in minimal, growth factor-free media, we generated neurons from human pluripotent stem cells in six days. Consistent with prior NGN2 studies, studies(7– 13) this protocol efficiently produced TUJ1^+^ neurons that expressed markers for excitatory cortical neurons and exhibited spontaneous, synchronous network activity by Day 14. While previous studies using CRISPR or lentivirus gene editing methods achieved 5–6% proliferative cells (7,33) or 100% neuronal purity (9), anti-mitotic agents were applied post-differentiation to eliminate non-neuronal cells. Our neuronal purity may be lower than other reports, but could similarly be enhanced with using antimitotic agents such as Ara-C or mitomycin C. Reducing factors in the induction media also lowers both complexity and cost while still yielding functionally mature neurons comparable to those generated under supplemented conditions (7,9,10). It is important to note that this protocol uses growth factor–supplemented media during the maturation phase to support the development of mature, electrically active neurons. For groups seeking to enhance maturation without reintroducing expensive neurotrophins, alternative low-cost strategies such as electrical stimulation (36), astrocyte co-culture (12,37,38), or the use of astrocyte-conditioned media (39) also promote functional neuronal development. Using chemically defined, xeno-free E8 medium further reduces hPSC costs relative to mTeSR or feeder-based systems, and expenses can be decreased even more by preparing the low-cost B8 formulation (40). Altogether, this workflow and potential adaptations provide a simple route to human neuron generation, balancing reduced media complexity with a modest purity trade-off, and enabling broader adoption of hiPSC-derived neuronal models across laboratories.

An additional advantage of this framework is its reliance on the PiggyBac transposon system, which provides a more accessible alternative to lentiviral delivery (9,13,16). High-titer lentiviral production requires specialized facilities, is time-intensive (41), often yields variable transduction efficiencies across cell lines (42), necessitating post-induction puromycin selection and contributing to inconsistent neuronal yields (9). In contrast, PiggyBac enables stable, high-capacity genomic integration without viral packaging, allowing transgene-positive hPSCs to be purified in a single puromycin step prior to differentiation, improving both yield and reproducibility. Ready-to-use PiggyBac vectors, including those for generating motor neurons and astrocytes, are increasingly available through repositories such as Addgene and the iNDi resource. As demonstrated here, secondary reporters can also be introduced for real-time tracking of neuronal morphology, expanding its utility for diverse downstream applications. Although PiggyBac integrates randomly and may introduce some variability in reprogramming efficiency, its simplicity, accessibility, and consistency across hPSC lines make it a practical tool that can be implemented in any lab with basic bacterial and molecular biology capabilities.

Overall, this simplified, low-cost protocol offers a more accessible human alternative to rodent primary neuron preparations, enabling both editing and differentiation of hiPSCs while requiring fewer specialized media components and lowering barriers for laboratories new to human neuron culture. Importantly, this workflow is suitable for editing and differentiating large patient-derived hiPSC collections, allowing rapid generation of neurons across diverse genetic backgrounds. Its use in KOLF2.1J hiPSCs is particularly valuable for disease modeling, as this line serves as the parental background for numerous mutations associated with neurodegenerative diseases through the iNDI (19). By facilitating broader adoption of human cell–based models, this approach aligns with the goals of the FDA Modernization Act 2.0 to reduce reliance on animal research.

## Statements

### Statement of Ethics

This study complies with the ISSCR Guidelines for the Conduct of Human Embryonic Stem Cell Research.

## Author Contributions

**N.A.P**. – Conceptualization, Investigation, Visualization, Writing - Original draft preparation, Formal analysis, Project administration. **O.E.K**. – Formal analysis, Investigation. **Z.L, F.S.S, K.E.M** – Investigation. **N.I**. – Conceptualization, Writing - Original draft preparation, Writing - Reviewing and editing, Supervision, Project administration, Funding acquisition.

## Data Availability Statement

The data supporting this research is archived and publicly available in the Tufts Dataverse repository (part of the Harvard Dataverse platform) and can be accessed via the following persistent identifier: *TBD*

## References

1. Hodge RD, Bakken TE, Miller JA, Smith KA, Barkan ER, Graybuck LT, et al. Conserved cell types with divergent features in human versus mouse cortex. Nature. 2019 Sept;573(7772):61–8.

2. Bakken TE, Jorstad NL, Hu Q, Lake BB, Tian W, Kalmbach BE, et al. Comparative cellular analysis of motor cortex in human, marmoset and mouse. Nature. 2021;598(7879):111–9.

3. Okano H, Morimoto S. iPSC-based disease modeling and drug discovery in cardinal neurodegenerative disorders. Cell Stem Cell. 2022 Feb 3;29(2):189–208.

4. Wu H, Xu J, Pang ZP, Ge W, Kim KJ, Blanchi B, et al. Integrative genomic and functional analyses reveal neuronal subtype differentiation bias in human embryonic stem cell lines. Proc Natl Acad Sci U S A. 2007 Aug 21;104(34):13821–6.

5. Strano A, Tuck E, Stubbs VE, Livesey FJ. Variable Outcomes in Neural Differentiation of Human PSCs Arise from Intrinsic Differences in Developmental Signaling Pathways. Cell Rep. 2020 June 9;31(10):107732.

6. Ortmann D, Vallier L. Variability of human pluripotent stem cell lines. Curr Opin Genet Dev. 2017 Oct 1;46:179–85.

7. Shan X, Zhang A, Rezzonico MG, Tsai MC, Sanchez-Priego C, Zhang Y, et al. Fully defined NGN2 neuron protocol reveals diverse signatures of neuronal maturation. Cell Rep Methods. 2024 Sept;4(9):100858.

8. Servetti M, Caramia M, Parodi G, Loiacono F, Nano E, Biddau G, et al. Optimization of Transcription Factor-Driven Neuronal Differentiation from Human Induced Pluripotent Stem Cells for Disease Modelling and Drug Screening. Stem Cell Rev Rep. 2025;21(3):816–33.

9. Zhang Y, Pak C, Han Y, Ahlenius H, Zhang Z, Chanda S, et al. Rapid Single-Step Induction of Functional Neurons from Human Pluripotent Stem Cells. Neuron. 2013 June 5;78(5):785–98.

10. Hulme AJ, Maksour S, St-Clair Glover M, Miellet S, Dottori M. Making neurons, made easy: The use of Neurogenin-2 in neuronal differentiation. Stem Cell Rep. 2021 Dec 30;17(1):14–34.

11. Lin HC, He Z, Ebert S, Schörnig M, Santel M, Nikolova MT, et al. NGN2 induces diverse neuron types from human pluripotency. Stem Cell Rep. 2021 Sept 14;16(9):2118–27.

12. Shih PY, Kreir M, Kumar D, Seibt F, Pestana F, Schmid B, et al. Development of a fully human assay combining NGN2-inducible neurons co-cultured with iPSC-derived astrocytes amenable for electrophysiological studies. Stem Cell Res. 2021 July 1;54:102386.

13. Nehme R, Zuccaro E, Ghosh SD, Li C, Sherwood JL, Pietilainen O, et al. Combining NGN2 Programming with Developmental Patterning Generates Human Excitatory Neurons with NMDAR-Mediated Synaptic Transmission. Cell Rep. 2018 May;23(8):2509–23.

14. Limone F, Guerra San Juan I, Mitchell JM, Smith JLM, Raghunathan K, Meyer D, et al. Efficient generation of lower induced motor neurons by coupling Ngn2 expression with developmental cues. Cell Rep. 2023 Jan;42(1):111896.

15. Wells MF, Nemesh J, Ghosh S, Mitchell JM, Salick MR, Mello CJ, et al. Natural variation in gene expression and viral susceptibility revealed by neural progenitor cell villages. Cell Stem Cell. 2023 Mar;30(3):312-332.e13.

16. Sheta R, Teixeira M, Idi W, Pierre M, De Rus Jacquet A, Emond V, et al. Combining NGN2 programming and dopaminergic patterning for a rapid and efficient generation of hiPSC-derived midbrain neurons. Sci Rep. 2022 Oct 13;12(1):17176.

17. Frega M, Van Gestel SHC, Linda K, Van Der Raadt J, Keller J, Van Rhijn JR, et al. Rapid Neuronal Differentiation of Induced Pluripotent Stem Cells for Measuring Network Activity on Micro-electrode Arrays. J Vis Exp. 2017 Jan 8;(119):54900.

18. Oh Y, Jang J. Directed Differentiation of Pluripotent Stem Cells by Transcription Factors. Mol Cells. 2019 Mar 1;42(3):200–9.

19. Pantazis CB, Yang A, Lara E, McDonough JA, Blauwendraat C, Peng L, et al. A reference human induced pluripotent stem cell line for large-scale collaborative studies. Cell Stem Cell. 2022 Dec 1;29(12):1685-1702.e22.

20. Ramos DM, Skarnes WC, Singleton AB, Cookson MR, Ward ME. Tackling neurodegenerative diseases with genomic engineering: A new stem cell initiative from the NIH. Neuron. 2021 Apr;109(7):1080–3.

21. Zhao S, Jiang E, Chen S, Gu Y, Shangguan AJ, Lv T, et al. PiggyBac transposon vectors: the tools of the human gene encoding. Transl Lung Cancer Res. 2016 Feb;5(1):120–5.

22. Lu X, Huang W. PiggyBac Mediated Multiplex Gene Transfer in Mouse Embryonic Stem Cell. PLOS ONE. 2014 Dec 17;9(12):e115072.

23. Gu J, Rollo B, Berecki G, Petrou S, Kwan P, Sumer H, et al. Generation of a stably transfected mouse embryonic stem cell line for inducible differentiation to excitatory neurons. Exp Cell Res. 2024 Feb 1;435(1):113902.

24. Lippmann ES, Estevez-Silva MC, Ashton RS. Defined Human Pluripotent Stem Cell Culture Enables Highly Efficient Neuroepithelium Derivation Without Small Molecule Inhibitors. Stem Cells. 2014 Apr 1;32(4):1032–42.

25. Chen G, Gulbranson DR, Hou Z, Bolin JM, Ruotti V, Probasco MD, et al. Chemically defined conditions for human iPSC derivation and culture. Nat Methods. 2011 May;8(5):424–9.

26. Chen Y, Tristan CA, Chen L, Jovanovic VM, Malley C, Chu PH, et al. A versatile polypharmacology platform promotes cytoprotection and viability of human pluripotent and differentiated cells. Nat Methods. 2021 May;18(5):528–41.

27. Cookson M. iNDI PiggyBac-TO-hNGN2 transfection protocol Version 1 v1 [Internet]. 2023 [cited 2025 Nov 13]. Available from: https://www.protocols.io/view/indi-piggybac-to-hngn2-transfection-protocol-versi-cyd6xs9e

28. Flores EL, Qi A, Reilly L, Santiana M, Not Provided Caroline P, Ward M, et al. iNDI Transcription Factor-NGN2 differentiation of human iPSC into cortical neurons Version 1 v1 [Internet]. 2021 [cited 2025 Nov 15]. Available from: https://www.protocols.io/view/indi-transcription-factor-ngn2-differentiation-of-b2whqfb6

29. Iyer NR, Shin J, Cuskey S, Tian Y, Nicol NR, Doersch TE, et al. Modular derivation of diverse, regionally discrete human posterior CNS neurons enables discovery of transcriptomic patterns. Sci Adv. 2022 Sept 30;8(39):eabn7430.

30. Han L, Mu S, He Z, Wang Z, Qu J, Ye W, et al. CNQX facilitates inhibitory synaptic transmission in rat hypoglossal nucleus. Brain Res. 2016 Apr 15;1637:71–80.

31. Li Q, Burrell BD. CNQX and AMPA inhibit electrical synaptic transmission: a potential interaction between electrical and glutamatergic synapses. Brain Res. 2008 Sept 4;1228:43–57.

32. Borghese L, Dolezalova D, Opitz T, Haupt S, Leinhaas A, Steinfarz B, et al. Inhibition of Notch Signaling in Human Embryonic Stem Cell–Derived Neural Stem Cells Delays G1/S Phase Transition and Accelerates Neuronal Differentiation In Vitro and In Vivo. Stem Cells. 2010 May 1;28(5):955–64.

33. Schörnig M, Ju X, Fast L, Ebert S, Weigert A, Kanton S, et al. Comparison of induced neurons reveals slower structural and functional maturation in humans than in apes. eLife. 2021 Jan 20;10:e59323.

34. Larsen KB, Lutterodt MC, Møllgård K, Møller M. Expression of the Homeobox Genes OTX2 and OTX1 in the Early Developing Human Brain. J Histochem Cytochem. 2010 July;58(7):669–78.

35. Nieto M, Monuki ES, Tang H, Imitola J, Haubst N, Khoury SJ, et al. Expression of Cux-1 and Cux-2 in the subventricular zone and upper layers II–IV of the cerebral cortex. J Comp Neurol. 2004 Nov 8;479(2):168–80.

36. Diego-Santiago MDP, González MU, Zamora Sánchez EM, Cortes-Carrillo N, Dotti C, Guix FX, et al. Bioelectric stimulation outperforms brain derived neurotrophic factor in promoting neuronal maturation. Sci Rep. 2025 Feb 8;15(1):4772.

37. Hedegaard A, Monzón-Sandoval J, Newey SE, Whiteley ES, Webber C, Akerman CJ. Promaturational Effects of Human iPSC-Derived Cortical Astrocytes upon iPSC-Derived Cortical Neurons. Stem Cell Rep. 2020 July;15(1):38–51.

38. Kuijlaars J, Oyelami T, Diels A, Rohrbacher J, Versweyveld S, Meneghello G, et al. Sustained synchronized neuronal network activity in a human astrocyte co-culture system. Sci Rep. 2016 Nov 7;6(1):36529.

39. Zheng H, Feng Y, Tang J, Yu F, Wang Z, Xu J, et al. Astrocyte-secreted cues promote neural maturation and augment activity in human forebrain organoids. Nat Commun. 2025 Mar 23;16(1):2845.

40. Kuo HH, Gao X, DeKeyser JM, Fetterman KA, Pinheiro EA, Weddle CJ, et al. Negligible-Cost and Weekend-Free Chemically Defined Human iPSC Culture. Stem Cell Rep. 2020 Feb;14(2):256–70.

41. Fernandopulle MS, Prestil R, Grunseich C, Wang C, Gan L, Ward ME. Transcription Factor– Mediated Differentiation of Human iPSCs into Neurons. Curr Protoc Cell Biol. 2018 June;79(1):e51.

42. Rapti K, Stillitano F, Karakikes I, Nonnenmacher M, Weber T, Hulot JS, et al. Effectiveness of gene delivery systems for pluripotent and differentiated cells. Mol Ther - Methods Clin Dev. 2015;2:14067.

