## Supplemental Table and Figures for "Rapid, Growth Factor-Reduced Differentiation of Functional Neurons from hiPSCs"

**Supplementary Table 1.** TaqMan assay information.

| Target Gene | TaqMan Assay ID |
|---|---|
| ***Oct3/4*** | Hs04260367_gH |
| ***Ngn2*** | Hs00702774_s1 |
| ***Pax6*** | Hs01088114_m1 |
| ***Otx2*** | Hs00222238_m1 |
| ***Cux2*** | Hs00322624_m1 |

**Supplementary Table 2.** Primary antibody manufacturer and dilution information.

| Antibody | Species | Manufacturer | Catalog # | Dilution |
|---|---|---|---|---|
| **OCT3/4** | Mouse | Santa Cruz Biotechnology | sc-5279 | 1:100 |
| **PAX6** | Rabbit | Biolegend | 901301 | 1:200 |
| **SOX2** | Goat | Abcam | ab239218 | 1:200 |
| **TUJ1** | Mouse | Biolegend | 801202 | 1:200 |
| **TUJ1** | Rabbit | Biolegend | 802001 | 1:200 |
| **MAP2** | Mouse | Millipore | MAB3418 | 1;500 |
| **VLGUT2** | Guinea Pig | Millipore | AB2251-I | 1:1000 |
| **SYN1** | Rabbit | Millipore | | 1:200 |

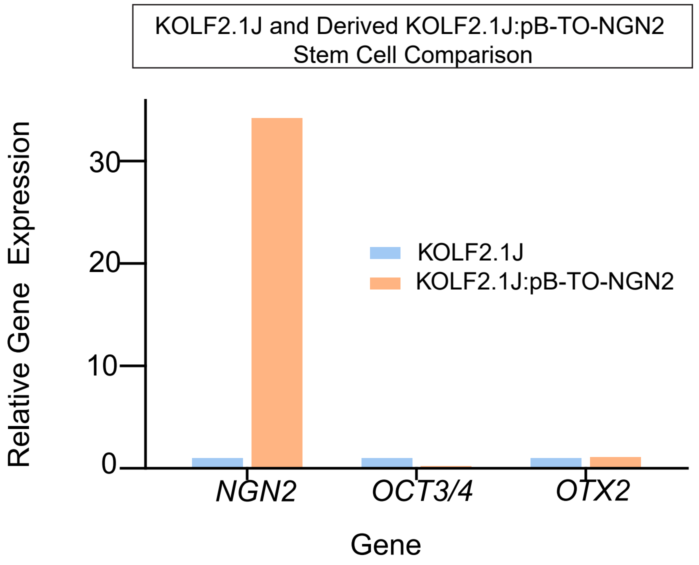


**Supplementary Figure 1. *NGN2* has higher gene expression in transfected stem cells.** qRT-PCR of *NGN2* (neuronal), *OCT3/4* (pluripotency), and *OTX2* (cortical forebrain) in KOLF2.1J:pB-TO-NGN2 hiPSCs compared to KOLF2.1J hiPSCs. (n=1)

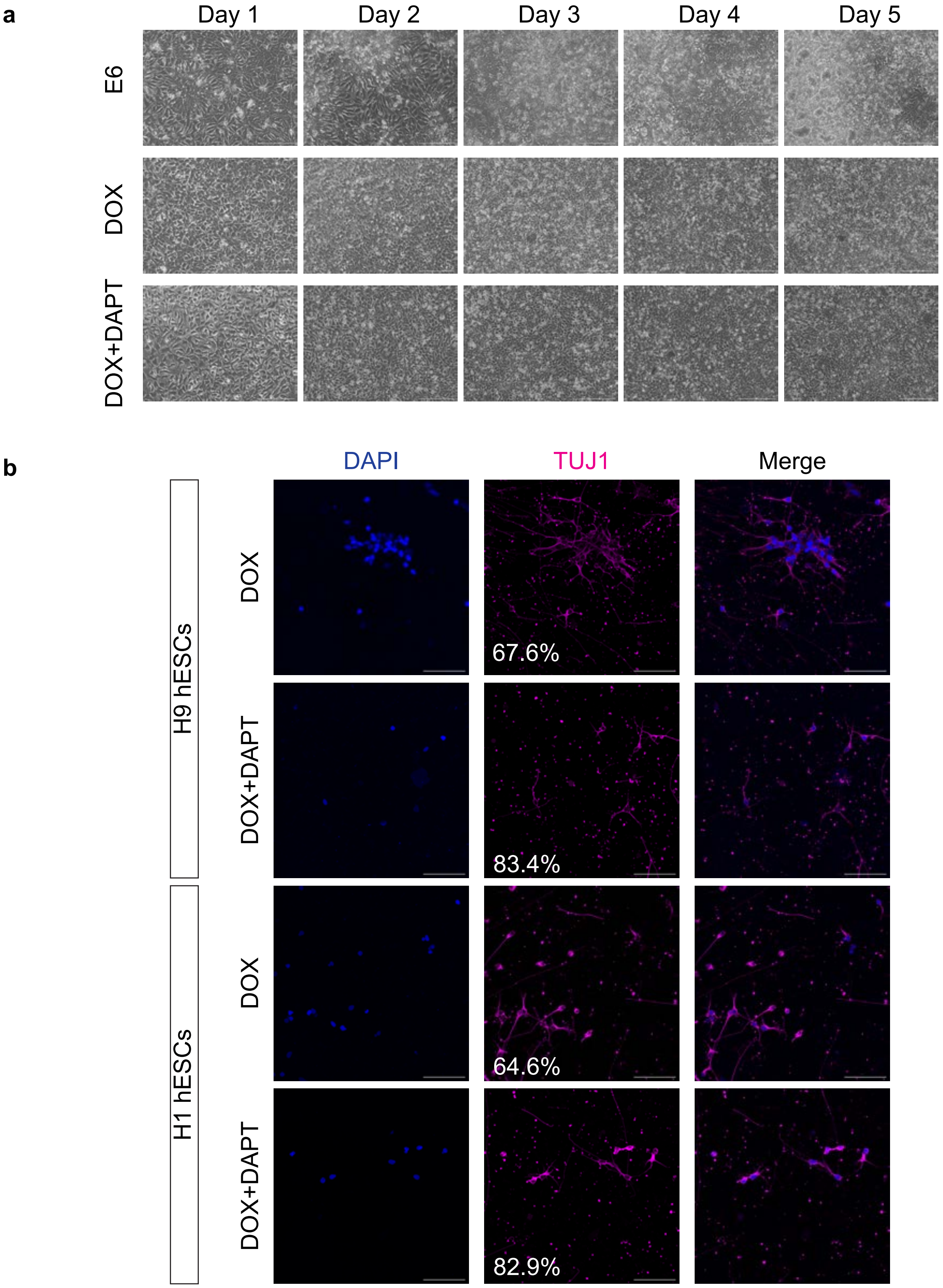
a
Day 1
Day 2
Day 3
Day 4
Day 5
E6
DOX
DOX+DAPT
b
DAPI
TUJ1
Merge
H9 hESCs
H1 hESCs
DOX
DOX+DAPT
67.6%
83.4%
64.6%
82.9%

**Supplementary Figure 2. Induction of PB-TO-NGN2 in KOLF2.1J, H1 and H9 hPSC**

**(a)** Phase contrast photos of NGN2 induction. E6, DOX, and DOX+DAPT conditions. Scale=150um

**(b)** Representative immunostaining images of quantification of neuronal conversion in H1:pB-TO-NGN2 and H9:pB-TO-NGN2 hPSC lines. Neurons were reseeded on Day 6 for staining. Percentages represent quantification of %TUJ1$^{+}$/DAPI$^{+}$ cells (n=1). Scale=75 μm.

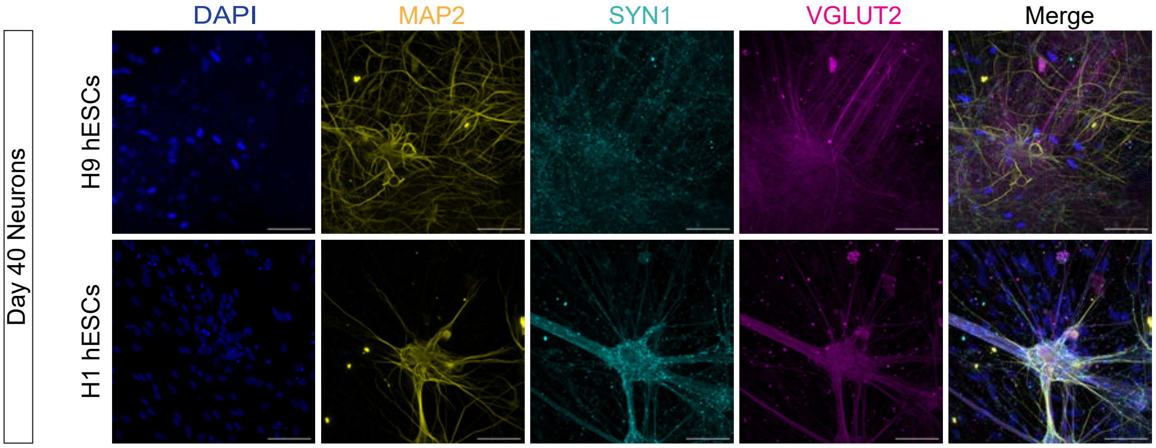


**Supplementary Figure 3. Day 40 H9 and H1 PB-TO-NGN2 Induced Neurons Express Mature Cortical Glutamatergic Markers**

Immunostaining of Day 40 H9:pB-TO-NGN2 and H1:pB-TO-NGN2 neurons for MAP2 (mature neurons), VGLUT2 (glutamatergic neurons), and SYN1 (pre-synaptic marker). Scale=75 μm.

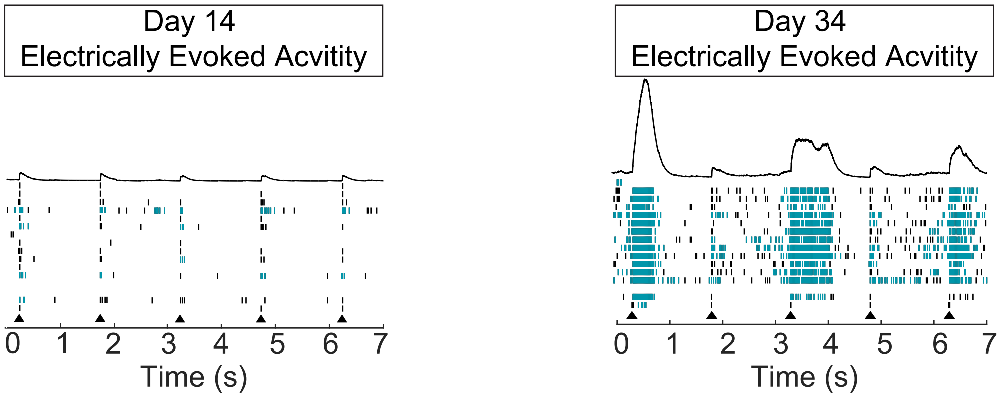


**Supplementary Figure 4. NGN2 induced neurons are responsive to electrical stimulus**

Representative raster plots from MEA recordings on Days 14 and 34 of culture demonstrating electrophysiological response to electrical stimulus.

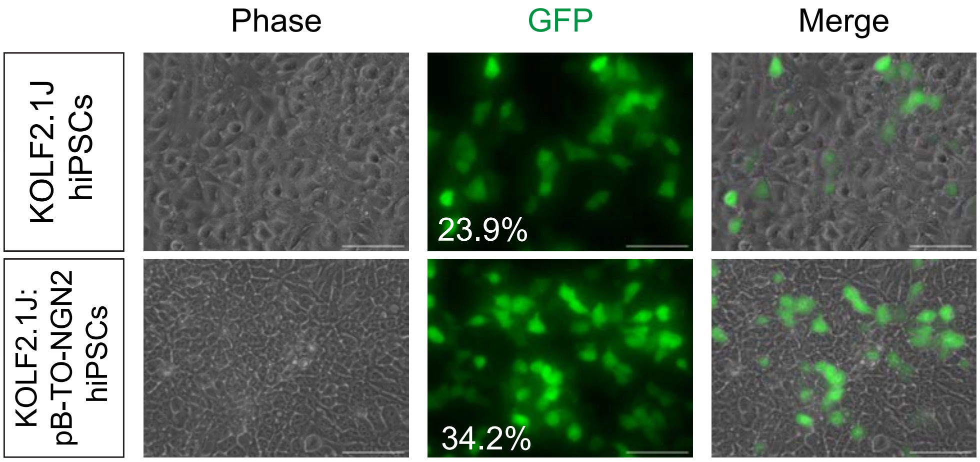


**Supplementary Figure 5. Secondary PiggyBac transfection of PB-CAG-eGFP.**
Percentage GFP expression 24 hours after transfection with PB-CAG-eGFP in KOLF2.1J and KOLF2.1J::pB-TO-NGN2 hiPSC lines. Scale =75 µm.
